# Astroglia in lateral habenula is essential for antidepressant efficacy of light

**DOI:** 10.1101/2020.07.31.230201

**Authors:** Sarah Delcourte, Adeline Etievant, Renaud Rovera, Damien Mor, Howard M Cooper, Hiep D. Le, April E. Williams, Satchidananda Panda, Rihab Azmani, Olivier Raineteau, Christine Coutanson, Ouria Dkhissi-Benyahya, Nasser Haddjeri

## Abstract

Successful antidepressant (AD) treatments are still difficult to achieve. Recently, bright light stimulation (BLS) was shown effective in non-seasonal depression but its mode of action remains elusive. We demonstrate here, using a new mouse model of depression resistant to ADs including ketamine, that chemogenetic activation of lateral habenula (LHb) astroglia prevented the potentiating effect of BLS on the AD response. Additionally, the beneficial action of BLS was associated with upregulation of a specific part of the prefrontal cortex opioid system. These results show that improved behavioral outcome produced by BLS requires habenular astroglia and endogenous opioids as crucial buffer systems.

## Introduction

Although major depression (MD) is the most common psychiatric disorder worldwide, fully effective treatments to cure MD are still awaiting. One of the safer, low-cost and non-invasive therapeutic candidates is bright light stimulation (BLS). Light has a critical broad role in health, acting directly or through the circadian system to modulate brain structures involved in sleep regulation, mood and cognition (**LeGates et al., 2014**). Hence, aberrant light cycles produce depressive behaviors and impair cognition in animals and in humans (**LeGates et al., 2014; Fernandez et al., 2018**). Moreover, the prevalence of MD is increased in night-workers, in whom both light exposure and circadian rhythmicity are altered. Inversely, depressed patients frequently display disturbed circadian rhythms and sleep/wake cycles (**McClung, 2013**). Since 3 decades, BLS is the first-line choice for the treatment of seasonal depression (**Wirz-Justice et al., 2004**). Surprisingly, BLS treatment has also proven to be effective in non-seasonal depression and even more efficient than the classical AD Prozac (**Lam et al., 2016**). Paradoxically, the mechanisms involved in the therapeutic effect of light are still unknown. The main objective of this study was to assess the involvement of the circadian, habenular and 5-HT systems, three important interactive brain networks regulating mental health (**Metzger et al., 2017**) and playing a key role in mood regulation by light (**LeGates et al., 2014; Fernandez et al., 2018; Huang et al., 2019**).

## Results & Discussion

The current ‘gold-standard’ behavioral test to examine the efficacy of potential AD agents is the forced swimming test (FST) in which ADs are well known to decrease immobility and increase active coping behavior (swimming and climbing) (**Porsolt et al., 1997**). To provide a reliable mice model of depression, we recently proposed that the timing of stress and testing is a critical factor for 5 days Repeated Forced Swim Stress (5d-RFSS) model (**Sun et al., 2015; Delcourte et al., 2017**). In the present study, 6-week old C57BL/6J mice socially isolated were forced to swim during the dark/active phase on 5 successive days for 10 minutes at zeitgeber time 14 (ZT14) and were then tested every week. A stress at ZT14 was extremely effective to induce a long-lasting (up to 8 weeks) depressive-like behavior (Fig. 1A). Therefore, mice exposed to the 5d-RFSS expressed a reduced sucrose preference indicating the presence of an anhedonia phenotype (Fig. 1B). In addition, the stressed mice exhibited higher anxiety and fear levels, measured as reduced time spent in the open arms in the elevated plus maze, as well as an increased latency to feed in the novelty suppressed feeding test (Fig. 1C, 1D). Treatment with the selective 5-HT reuptake inhibitor AD escitalopram, or the NMDA receptor antagonist ketamine produced an AD-like behavior in naïve mice (Fig 1F-H). In contrast, these treatments were ineffective to reverse the behavioral despair in 5d-RFSS mice (Fig. 1E-G), thus revealing 5d-RFSS as a new pharmaco-resistant model of depression. Remarkably, among all antidepressants tested in the current study, only a sub-chronic treatment with the mixed opioid agent buprenorphine (a potent mu opioid receptor partial agonist and kappa opioid receptor antagonist, **Falcon et al., 2016**) showed an AD-like response in 5d-RFSS mice (Fig S1).

**Figure 1.**
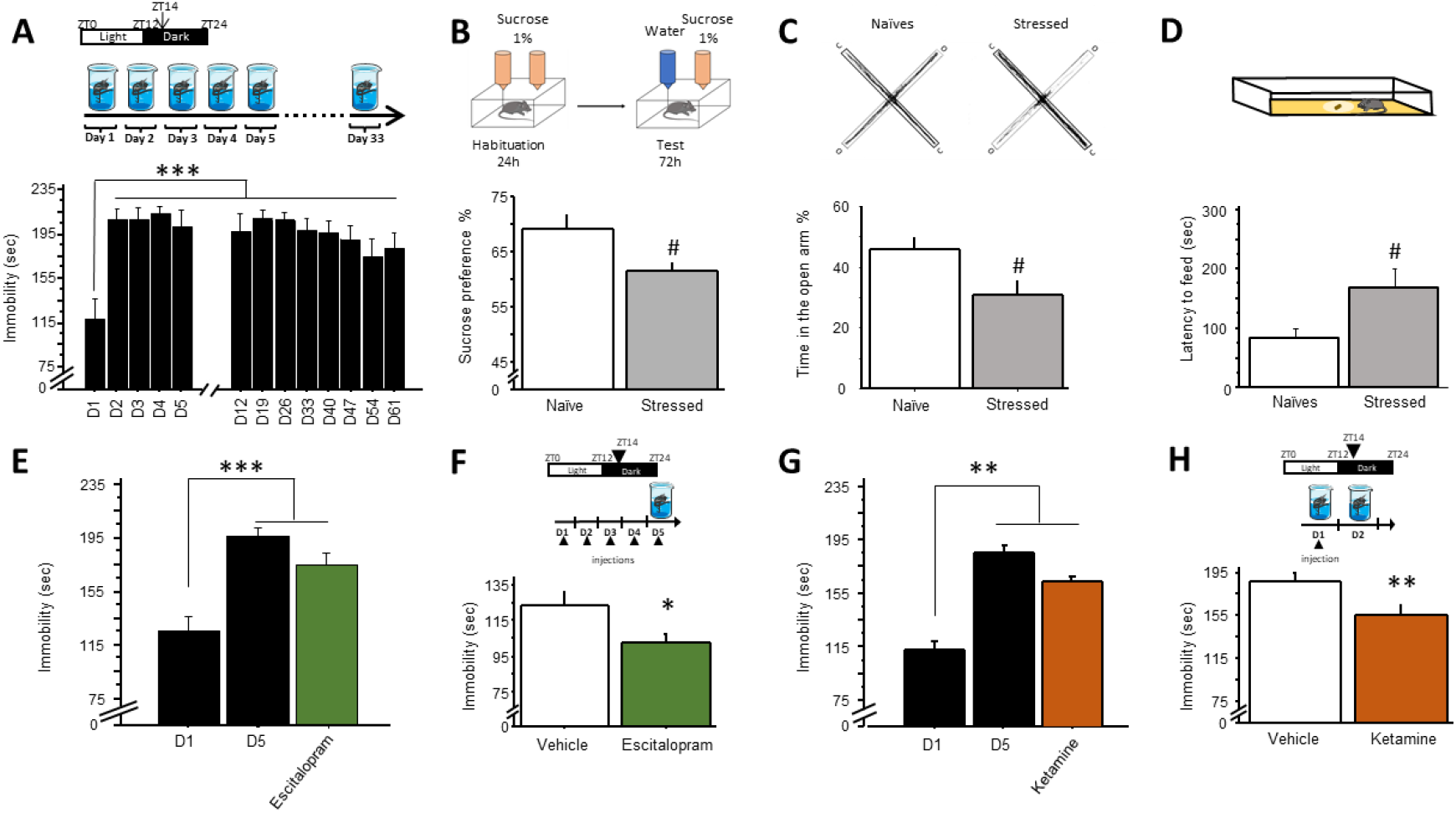
5d-RFSS: a resistant model of depression. **(A)** *5d-RFSS*: 6 weeks-old C57BL/6J mice were forced to swim for 5 successive days [D] and for 10 minutes at ZT14. Animals were then tested every week (n=8), ***p<0.0001 vs D1. **(B)** *Sucrose Preference Test*, 48h post stress (n=8), ## p<0.001. **(C)** *Elevated plus maze*, 1 week post-stress (n=8-13). C=closed arm, O=open arm, # p<0.05. **(D)** *Novelty suppressed feeding test* (n=9-12), # p<0.05. **(E)** *Effect of a sub-chronic escitalopram treatment* (10 mg/kg i.p) 5 consecutive days from D10 to D15 treatment, (n=8) p<0.0001 vs D1. **(F)** *Effect of a sub chronic escitalopram treatment on naïve mice* (n=8) * p<0.05 **(G)** *Effect of acute ketamine treatment* (10mg/kg, i.p), n=6, **p<0.001 vs D1. **(H)** *Effect of acute ketamine treatment on naïve mice* (10mg/kg, i.p) unpaired Student’s t-tests; ** p<0.01 (n=13-15). Data are expressed as means ± S.E.M.

Using the FST as a main AD readout, we then assessed the effect of BLS alone or in combination with AD agents on this refractory model. BLS failed to induce an AD response given alone (Fig. 2B). While a combination of sub-effective doses of ketamine and the muscarinic receptor antagonist scopolamine produced an AD-like behavior in naïve mice (**Petryshen et al., 2016**; Fig.2A), the latter antagonists did not reverse the behavioral despair induced by the 5d-RFSS (Fig. 2C). Importantly, we found that BLS added to a combination of ketamine and scopolamine, reduced the immobility time observed in the FST, indicating that BLS potentiated the AD response of the latter AD treatment (Fig. 2C).

**Figure 2.**
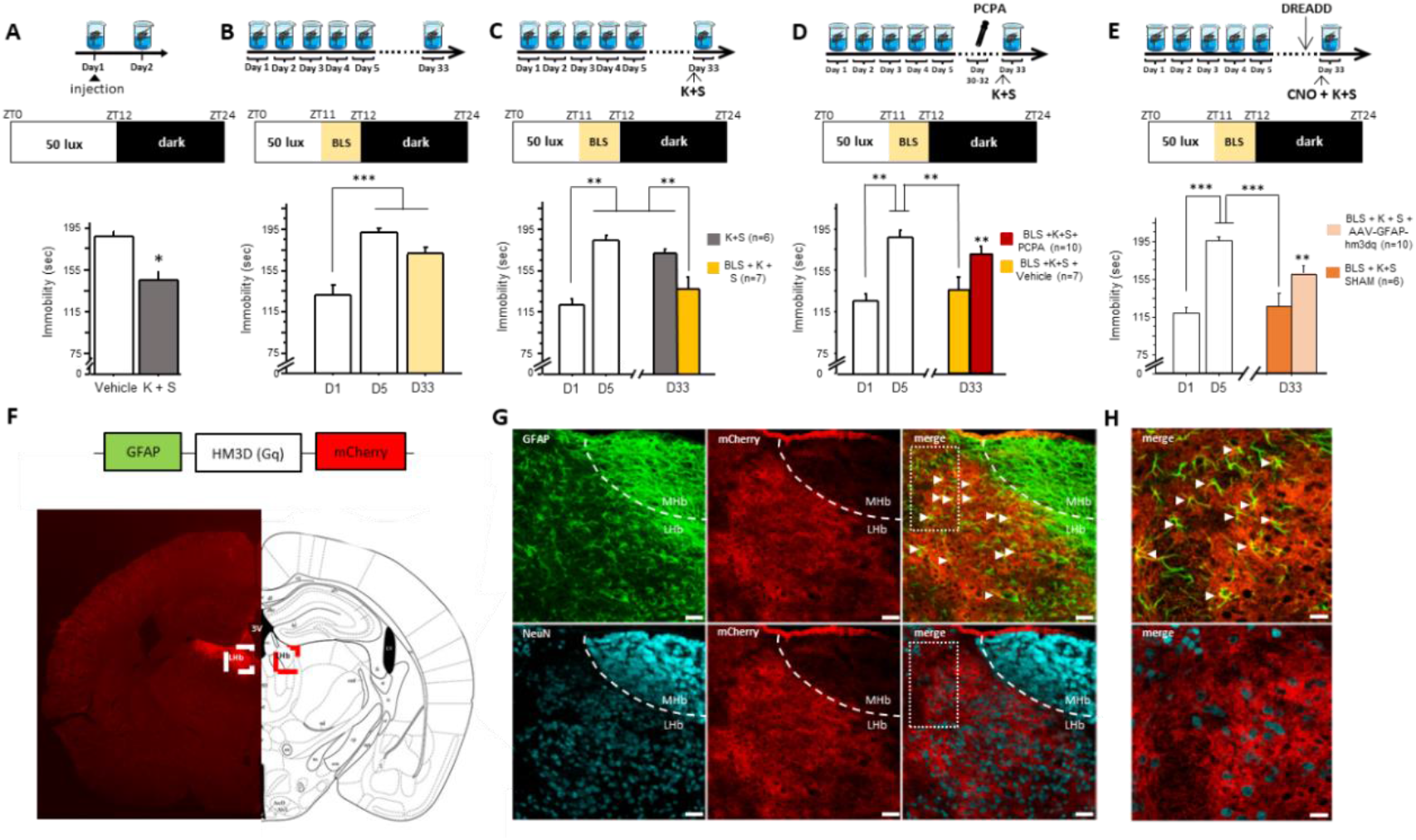
BLS potentiates the AD response of a combination of ketamine and scopolamine. **(A)** *Effect of a pharmacological co-treatment with Ketamine [K] (3mg/kg, i.p.) and Scopolamine [S]* (0.1 mg/kg, i.p.) in naïve mice (* P<0.05, n=8). **(B)** *Effect of a BLS* on the increase of immobility time in 5d-RFSS model (n= 6), *** P<0.0001 vs D1. **(C)** *Effect of a combination of BLS and pharmacological co-treatment* with K + S (n=6-7). ** P<0.01 vs D1 or D33. On day 33, the two groups were compared. **(D)** *Involvement of 5-HT neurotransmission*: mice received a dose of 150 mg/kg/day (i.p) of PCPA to reduce the levels of 5-HT (n= 7-10). ** P<0.01 vs D1 or D33. On day 33, the two groups were compared. **(E)** *Implication of LHb astrocytes* in the AD effect of BLS: Following 5d-RFSS, mice were intracerebrally injected with vehicle or a AAV-GFAP-Gq-m-cherry virus in the LHb. A 4-week BLS was then applied. At D33, mice received an injection of Clozapine-n-oxide [CNO] (1 mg/kg, i.p.) 1 hour before the FST. Half an hour after the CNO injection, co-treatment [K+S] (respectively 3 and 0.1 mg/kg i.p) was administered to the mice (n=6-10). *** P<0.01 vs D1 or D33. On day 33, the two groups were compared. **(F)** *Localization of AAV-GFAP-Gq-m-cherry virus in the LHb*: Coronal image of brain section showing mCherry labelling (red) in the LHb (white square). **(G,H)** *Overview of astroglial and neuronal staining in the LHb*: Anti-GFAP (green) and anti-NeuN (blue). Scale bars = 150μm (G), 75μm (H). Datas are expressed as means ± S.E.M.

Several lines of evidence significantly link the effects of light to 5-HT regulation of mood (**Gonzalez & Aston-Jones, 2008; Defrancesco et al., 2013**). Interestingly, the AD-like effect of ketamine, but not scopolamine, was abolished by 5-HT depletion produced by the tryptophan hydroxylase inhibitor, para-chlorophenylalanine (PCPA) (**Giglucci et al., 2013; Palucha-Poniewiera et al., 2017**). Accordingly, our results showed that the potentiating effect of BLS on the action of the ketamine/scopolamine combination was prevented by PCPA pre-treatment, revealing a permissive role of the 5-HT system in the AD behavioral phenotype (Fig. 2D).

LHb has emerged as a key structure interface between light effects and 5-HT regulation of mood. LHb, which receives projections from the intrinsically photosensitive retinal ganglion cells (**Hattar et al., 2006**), is anatomically and functionally connected to the raphe 5-HT nuclei (**Pasquier et al., 1976**). While it was firstly shown that pseudo-depressed mice display increased activity of LHb neurons (**Lecca et al., 2016**), **Yang et al (2018)** recently reported in the LHb that coordinated activity of NMDA receptors and T-type voltage-sensitive Ca2+ channels cause neuron firing to occur in a pattern of rapid bursts and that such aberrant firing leads to depressive-like symptoms. Additionally, **Cui et al. (2018)** revealed the involvement of LHb astroglia in the increases of burst firing mode and depressive-like phenotype. Hence, we sought to determine whether selective chemogenetic activation of transduced astroglia in LHb, could counteract the potentiating AD action of BLS in 5d-RFSS mice. Injection of the GFAP-Gq-DREADD virus in the LHb showed fields of mCherry-positive cells, that consistently express the GFAP astroglia marker and lack the NeuN neuronal marker (Fig. 2G-H). Remarkably, clozapine-n-oxide (CNO) administration, by producing a selective and robust activation of astrocytes in mice expressing Gq-DREADD in LHb astrocytes, significantly prevented the potentiating effect of BLS on the action of the ketamine/scopolamine combination (Fig. 2E). This result reveals a crucial involvement of LHb astroglia as well as a permissive role of 5-HT and opioid systems in the beneficial action of BLS in the 5d-RFSS pharmaco-resistant model. Recently, the antidepressant effects of ketamine were shown to be attenuated by opioid receptor blockade in both human and rodent (**Williams et al., 2019; Klein et al., 2020**). Importantly, the reducing action of ketamine on LHb cellular hyperactivity is also prevented by opioid receptor blockade (**Klein et al., 2020**). To further characterize both 5d-RFSS model and BLS action, we performed RNA-seq in the PFC, a key brain region involved in both MD (**Krishnan & Nestler, 2008**) and antidepressant response (including ketamine) (**Gerhard et al., 2019**). Remarkably and in full agreement with the AD response of buprenorphine, the efficient co-treatment with light modified opioid system gene expression in PFC with a peculiar enhancement of Opioid Receptor Mu-type 1 (OPRM1), Opioid Receptor Kappa-type 1 (OPRK1), Proenkephalin (PENK) and Prodynorphin (PDYN) gene expressions, thus demonstrating for the first time that BLS potentiated the effect of pharmacological treatments by mechanisms involving the expression of opioid-related genes in PFC (Fig S2).

As a translational outcome, the current study will certainly have far-reaching impact on our understanding of the mechanisms involved in light effects on depression to ultimately optimize therapeutic strategies in drug-resistant or refractory depressed patients (including partial-responders to ketamine) and proposes BLS as a beneficial strategy against the possible deleterious effect induced by social isolation such as e.g. the current one required against the Covid-19’s pandemic.

## Methods

### Animals

All animal procedures were in strict accordance with current national and international regulations on animal care, housing, breeding, and experimentation and were approved by the regional ethics committee CELYNE (C2EA42-13-02-0402-005). All efforts were made to minimize suffering. 6 week-old C57BL/6J male mice (25–30 g on arrival; Charles River Laboratories) were individually housed in a temperature- and humidity-controlled environment with a 12 h dark/12 light (12L/12D) cycle. Mice received food and water ad libitum and were allowed 2 weeks to acclimate before the experiments.

### 5 days Repeated Forced Swim Stress (5d-RFSS)

Two hours after lights off (ZT14), mice were placed in a tank filled with water (25°C) and forced to swim 10 minutes daily for 5 consecutive days. The immobility time of the first 4 minutes of the stress was analyzed. Then, every week, a forced swim test (FST) was realized to follow the evolution of the immobility time.

### Sucrose preference test

Mice were habituated to 2 bottles of 1% sucrose during 24 h. Then, they were given the choice to drink from 2 bottles (1% sucrose solution and tap water bottle) during 72 h. The positions of the bottles in the cage were switched every 24 h to avoid possible side-preference effects.

### Elevated plus maze

Animals were placed in the room half an hour before the beginning of the test. Experiments were realized during the light phase of the 12L/12D cycle. The apparatus consisted of 2 Plexiglas open arms (6×96.5 cm), and 2 closed arms (6×95.5 cm), surrounded by 15cm black high walls elevated 50 cm above the floor. At the beginning of the test, the mouse was placed in the central platform of the maze, facing an open arm. Each session lasted 6 min and was video recorded to analyze the time spent in the open arms.

### Novelty suppressed feeding test

The latency to begin eating a pellet of food in a frightening environment is considered as a measure of anxio/depressive-like behavior. Animals were food-deprived for 24h prior to behavioral testing. Mice were individually placed in a brightly illuminated plastic box (50 cm ×40 cm × 20 cm) covered with wooden bedding. A single food pellet (regular chow) was placed on a white paper platform positioned in the center of the arena. The latency to eat (defined as the mouse sitting on its haunches and biting the pellet with the use of forepaws) was timed. The NSF test was carried out during a 5 min period.

### Drugs

Escitalopram was supplied by H. Lundbeck A/S. Ketamine hydrochloride was purchased from Sigma-Aldrich. Scopolamine hydrobromide and Clozapine-N-oxide (CNO) were purchased from Tocris Biosciences, Buprenorphine (Subutex) was kindly provided by Bruno Guiard (CRCA, Toulouse). All drugs were dissolved in a 0,9% saline solution. Escitalopram sub-chronic treatment was administered at 10mg/kg, i.p during 5 consecutive days following the 5d-RFSS, or during the five preceding the FST for the naïve group. Buprenorphine (0,3 mg/kg, i.p.) was administered every two days five times. Ketamine hydrochloride was administered acutely at 10 mg/kg alone or in combination with scopolamine (0,1 mg/kg, i.p), at 3 mg/kg, i.p. Ketamine and scopolamine were administered 30 minutes before the forced swimming test. Clozapine-N-oxide (CNO) was dissolved in saline and administered at 1mg/kg, i.p. 30 minutes before the FST.

### Light stimulation protocols

At the end of the 5d-RFSS, mice were exposed every day to a bright white light (1000 lux; LED bulb) for one hour at ZT11. The effect of light exposure on depressive-like behavior was analyzed every week during 4 weeks post 5d-RFSS.

### PCPA treatment

Three days before the 4-week’s FST, animals were injected each day with 150 mg/kg of 4-Chloro-DL-phenylalanine (PCPA, Sigma Aldrich), the last day of injection was 24h before the last FST.

### Surgery

Mice were anesthetized with Chloral Hydrate (30m/kg, i.p.; SIGMA) and Xylazine (12 mg/kg i.p.; Bayer). There were implanted with cannulas for the virus microinjections. The vector, containing the virus, was dissolved in PBS + MgCl2 + KCl at a concentration of 6,0.1012 vg/mL. 0,2 μl of ssAAV-5/2-hGFAP-hM3D(Gq)-mCherry-WPRE-hGHp(A) (GFAP-Gq-DREADD; Viral vector Facility-Zurich University) was infused bilaterally into the LHb (AP + 4.24mm; ML± 0.48; DV + 4.2mm from bregma with a 45° angle) using 33 gauge injectors at rate of 0.05 μl/min.

### Immunohistochemical Staining

Animals were given pentobarbital (50 mg/kg, i.p.) and trans-cardiac perfused with 4% paraformaldehyde (PFA). Brains were removed and post-fixed in 4% PFA at 4°C for an additional 24 h, rinsed in PBS (phosphate-buffered saline, pH 7.4), and cryo-protected in 30% sucrose in 0.1 M PBS for an additional 48 h at 4°C. Brains were harvested and sliced at 30 μm, and incubated for 24 h with anti-GFAP (Mouse, 1:500, Sigma), anti-NeuN (Guinea Pig, 1:500, Synaptic systems) and anti RFP (Rabbit, 1:1000, MBL) antibodies in PBS + 5% normal goat serum. RFP signal was amplified using a biotinylated anti-rabbit secondary antibody (Goat, 1:200, Vector Biosystem) and streptavidin-DTAF complex (1:250, Jackson Labs Technologies) for 30 min. GFAP, NeuN, and RFP stainings were visualized using respectively Alexa Fluor 488, 647 (Donkey 1:500 Jackson Immunoresearch) and Streptavidin-Cy3 (1:500, Jackson Immunoresearch). Images were obtained with a Leica SP5 confocal (Leica Microsystems). Z-series images were taken at 2 μm intervals.

### RNA extraction

Two days after the last forced swim test, mice were euthanized. Both sides of the medial prefrontal cortices were dissected on ice and stored at −80°C. Total RNAs were extracted using Trizol reagent (Invitrogen) according to manufacturer’s instructions.

### Libraries preparation and high-throughput sequencing

Libraries were prepared using Illumina’s TruSeq Stranded mRNA HT kit according to manufacturer’s instructions. In brief, total RNA starting with 700ng was poly-A selected, fragmented by metal-ion hydrolysis and then converted to cDNA using SuperScript II. The cDNA was then end-repaired, adenylated and ligated with Illumina sequencing adapters. Finally, the libraries were enriched by 15 cycles of PCR amplification. Libraries were pooled and sequenced using an Illumina HiSeq 2500 with 50-bp single-read chemistry.

### Transcriptome data analysis

Whole-transcriptome profiling of prefrontal cortices was performed by RNASeq. Sequenced reads were mapped to the reference mouse genome using (org.Mm.eg.db, http://bioconductor.org/packages/release/data/annotation/html/org.Mm.eg.db.html). Read counts were generated using Homer and normalized counts were used in DESeq2 (Bioconductor, http://bioconductor.org/packages/release/bioc/html/DESeq2.html) to measure differential expression. The RNASeq raw data are available at GEO under accession number GSE143820. The candidate genes were screened from a comparison of control and stressed mice, as well as of stressed mice treated with light, scopolamine and ketamine (i.e. KScL mice) as follows: first, the normalized counts were cut-off at 25 in at least one of the samples, then the gene lists were generated based on adjusted p-value <0.01. Lists of genes were submitted to DAVID (https://david.ncifcrf.gov) for functional enrichment analysis.

## Author Contributions

S.D., A.E., O.D-B and N.H. designed research; S.D. and A.E. performed research; R.R., D.M. and C.C contributed to the behavioral experiments and immunochemistry; S.D., A.E., R. A., H.H.L., A. E., O.D-B and N.H. analyzed the data; H.M.C. S. P. and O.R. critically reviewed the manuscript. S.D., A.E., O.D-B and N.H. wrote the paper.

## Acknowledgments

This research was supported by “La Région Rhône-Alpes-SCUSI 2018-#R18119CC” and “l’INSERM” and the ANR Grant “progenID”. Whole transcriptome profiling was supported by NIH grant EY 016807.

---

**Table I:**
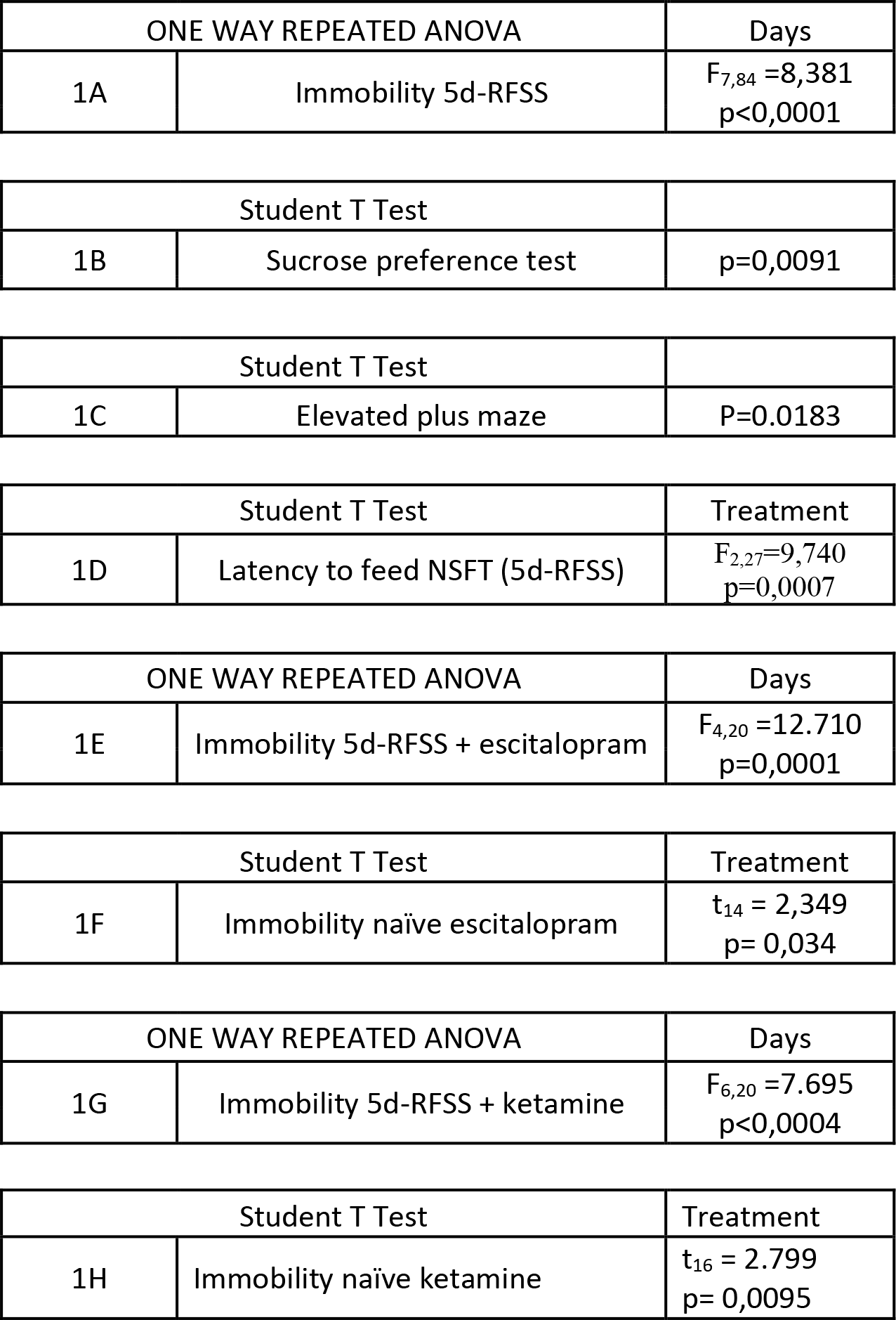
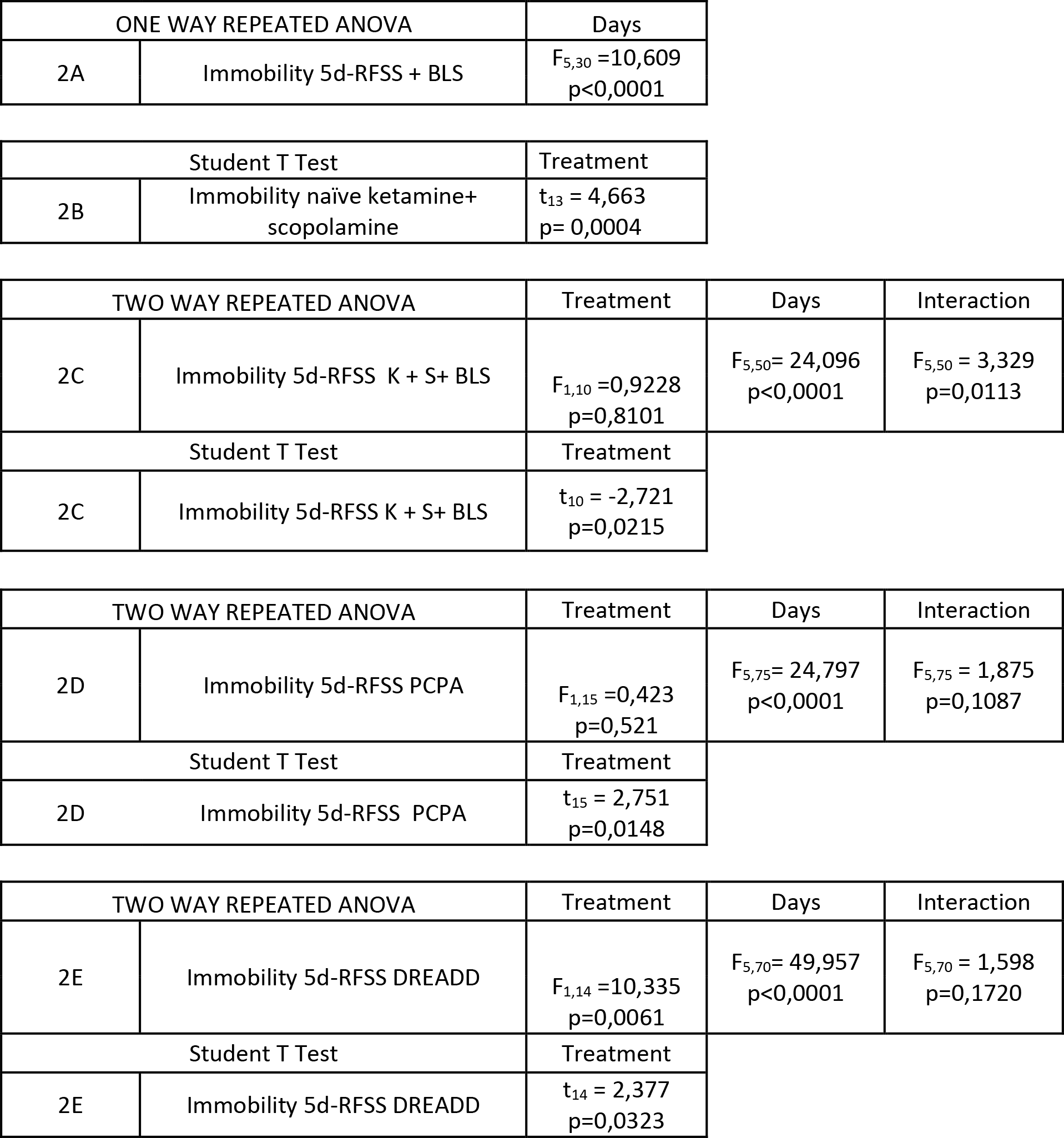
Statistical analysis

## References

Cui Y, Yang Y, Ni Z, Dong Y, Cai G, Foncelle A et al. Astroglial Kir4.1 in the lateral habenula drives neuronal bursts in depression. Nature. 2018; 554(7692): 323–327. doi:10.1038/nature25752

Defrancesco M, Niederstätter H, Parson W, Kemmler G, Hinterhuber H, Marksteiner J, Deisenhammer EA Bright.ambient light conditions reduce the effect of tryptophan depletion in healthy females. Psychiatry Research. 2013; 210(1): 109–14. doi: 10.1016/j.psychres.2013.02.008

Delcourte S, Dkhissi-Benyahya O, Cooper HM, & Haddjeri N. Stress Models of Depression: A Question of Bad Timing. ENeuro. 2017;4(2): 1–2. doi: 10.1523/ENEURO.0045-17.2017

Falcon E, Maier K, Robinson SA, Hill-Smith TE, Lucki I. Effects of buprenorphine on behavioral tests for antidepressant and anxiolytic drugs in mice. Psychopharmacology (Berl). 2015;232(5):907–915. doi: 10.1007/s00213-014-3723-y

Fernandez DC, Fogerson PM, Lazzerini Ospri L, Thomsen MB, Layne RM, Severin D et al. Light Affects Mood and Learning through Distinct Retina-Brain Pathways. Cell 2018; 175(1): 1–14. doi: 10.1016/j.cell.2018.08.004

Gerhard DM, Pothula S, Liu RJ, Wu M, Li XY, Girgenti MJ et al. GABA interneurons are the cellular trigger for ketamine’s rapid antidepressant actions. J Clin Invest. 2020; 130(3):1336–1349 doi: 10.1172/JCI130808

Gigliucci V, O’Dowd G, Casey S, Egan D, Gibney S, Harkin A. Ketamine elicits sustained antidepressant-like activity via a serotonin-dependent mechanism. Psychopharmacology. 2013; 228(1): 157–166. doi: 10.1007/s00213-013-3024-x

Gonzalez MMC., & Aston-Jones G. Light deprivation damages monoamine neurons and produces a depressive behavioral phenotype in rats. Proceedings of the National Academy of Sciences of the United States of America. 2008; 105(12): 4898–4903. doi: 10.1073/pnas.0703615105

Hattar S, Kumar M, Park A, & Tong P. Central Projections of Melanopsin-Expressing Retinal Ganglion Cells in the Mouse. Journal of Comparative Neurology. 2006; 497(3): 326–349. doi: 10.1002/cne.20970

Huang L, Xi Y, Peng Y, Yang Y, Huang X, Fu Y et al. A Visual Circuit Related to Habenula Underlies the Antidepressive Effects of Light Therapy. 2019; Neuron 102(1):128–142.e8. doi: 10.1016/j.neuron.2019.01.037

Klein ME, Chandra J, Sheriff S, Malinow R. Opioid system is necessary but not sufficient for antidepressive actions of ketamine in rodents. Proc Natl Acad Sci U S A. 2020;117(5):2656–2662. doi: 10.1073/pnas.1916570117

Krishnan V & Nestler EJ. The molecular neurobiology of depression. Nature. 2008; 455(7215): 894–902.

Lam RW, Levitt AJ, Levitan RD, Michalak EE, Cheung AH, Morehouse R et al. Efficacy of Bright Light Treatment, Fluoxetine, and the Combination in Patients With Nonseasonal Major Depressive Disorder. JAMA Psychiatry. 2016; 73(1): 56–63. doi: 10.1038/nature07455

Lecca S, Pelosi A, Tchenio A, Moutkine I, Lujan R, Hervé D, Mameli M. Rescue of GABAB and GIRK function in the lateral habenula by protein phosphatase 2A inhibition ameliorates depression-like phenotypes in mice. Nature Medicine. 2016; 22: 254–261. doi: 10.1038/nm.4037

LeGates TA, Fernandez DC, & Hattar S. Light as a central modulator of circadian rhythms, sleep and affect. Nat Rev Neurosci. 2014;18(9): 1199–1216. doi: 10.1038/nrn3743

McClung CA. How Might Circadian Rhythms Control Mood? Let Me Count the Ways… Biological Psychiatry. 2013; 74(4): 242–249. doi: 10.1016/j.biopsych.2013.02.019

Metzger M, Bueno D, and Lima LB. The lateral habenula and the serotonergic system. Pharmacology Biochemistry and Behavior. 2017; 162: 22–28. doi: 10.1016/j.pbb.2017.05.007

Pałucha-Poniewiera A, Podkowa K, Lenda T, & Pilc A. The involvement of monoaminergic neurotransmission in the antidepressant-like action of scopolamine in the tail suspension test. Progress in Neuro-Psychopharmacology and Biological Psychiatry. 2017; 79: 155–161. doi: 10.1016/j.pnpbp.2017.06.022

Pasquier DA, Anderson C, Forbes WB, & Morgane PJ. Horseradish peroxidase tracing of the lateral habenular-midbrain raphe nuclei connections in the rat. Brain Research Bulletin. 1976; 1(5): 443–451. doi: 10.1016/0361-9230(76)90114-3

Petryshen TL, Lewis MC, Dennehy KA., Garza JC., & Fava M. Antidepressant-like effect of low dose ketamine and scopolamine co-treatment in mice. Neuroscience Letters. 2016; 620: 70–73. doi: 10.1016/j.neulet.2016.03.051

Porsolt RD, Le Pichon M, Jalfre M. Depression: a new animal model sensitive to antidepressant treatments. Nature. 1977;21 266(5604):730–2. doi: 10.1038/266730a0

Sun P, Zhang Q, Zhang Y, Wang F, Chen R, Yamamoto R, Kato N. Homer1a-dependent recovery from depression-like behavior by photic stimulation in mice. Physiol Behav. 2015; 147:334–41. doi: 10.1016/j.physbeh.2015.05.007

Williams NR, Heifets BD, Bentzley BS, et al. Attenuation of antidepressant and antisuicidal effects of ketamine by opioid receptor antagonism. Mol Psychiatry. 2019;24(12):1779–1786. doi: 10.1038/s41380-019-0503-4

Wirz-Justice A, Terman M, Oren DA, Goodwin FK, Kripke DF, Whybrow PC et al. Brightening Depression. Science. 2004; 303: 467–468. doi: 10.1126/science.303.5657.467c

Yang Y, Cui Y, Sang K, Dong Y, Ni Z, Ma S, Hu H. Ketamine blocks bursting in the lateral habenula to rapidly relieve depression. Nature. 2018; 554(7692): 317–322. doi: 10.1038/nature25509

