## Supplemental Fig + Table for "Astroglia in lateral habenula is essential for antidepressant efficacy of light"

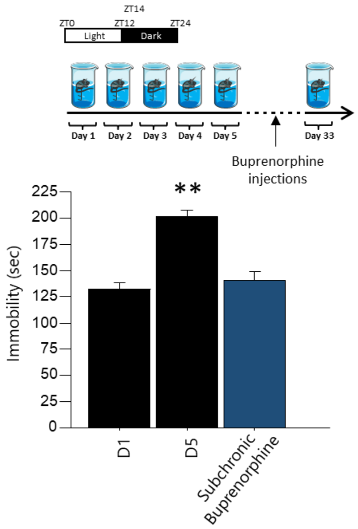

**Figure S1 : Effect of µ opioid receptor partial agonist buprenorphine on behavioral despair following 5d-RFSS**

Buprenorphine (0,3 mg/kg) was administered ever two days five times intraperitoneally before the final Forced swim test (n=8), One way ANOVA, ** p<0,01, Data are expressed as means ± S.E.M.

**Table S1**

Pairwise Intersections

| | stress vs stress KSC (4371) | stress vs stressL (6444) | stress vs stress KSCL (6884) | naives vs naives KSC (445) | genes opoids (20) |
|---|---|---|---|---|---|
| Naive vs stress (359) | 331 | 327 | 330 | 9 | 0 |
| stress vs stress KSC (4371) | | 4078 | 3994 | 72 | 5 |
| stress vs stressL (6444) | | | 5659 | 109 | 8 |
| stress vs stress KSCL (6884) | | | | 114 | 12 |
| naives vs naives KSC (445) | | | | | 0 |

*5 genes identity*: Gnas, Grk2, Ppp1r9b, Syp, Wls
*8 genes identity*: Gnas, Grk2, Ppp1r9b, Syp, Wls, Flna, Gpr88, Ppp1r1b
*12 genes identity*: Gnas, Grk2, Ppp1r9b, Syp, Wls, Flna, Gpr88, Ppp1r1b,
**Oprk1, Oprm1, Pdyn, Penk**

| **Table 2** | | | | | | |
|---|---|---|---|---|---|---|
| **Comparison Stressed vs Stressed KSCL** | | | | | | |
| | **baseMean** | **log2FoldChange** | **lfcSE** | **stat** | **pvalue** | **padj** |
| Penk | 2089,705233 | 1,675586683 | 0,124692 | 13,43782 | 3,63E-41 | 4,33E-38 |
| Oprk1 | 268,3821797 | 1,8540034 | 0,245623 | 7,548157 | 4,41E-14 | 1,63E-12 |
| Gpr88 | 1567,130465 | 2,416881132 | 0,139894 | 17,27653 | 7,07E-67 | 5,06E-63 |
| Pdyn | 500,8629163 | 1,159667041 | 0,159522 | 7,269625 | 3,60E-13 | 1,16E-11 |
| Oprm1 | 26,14590741 | 1,741472598 | 0,626467 | 2,779831 | 0,005439 | 0,015952 |
| Ppp1r1b | 1495,136606 | 1,071047983 | 0,128301 | 8,347916 | 6,95E-17 | 3,96E-15 |
| Wls | 982,843685 | 0,492962095 | 0,12616 | 3,907441 | 9,33E-05 | 0,000472 |
| *Positive values: upregulation in stressed KSCL group* | | | | | | |
| Ppp1r9b | 5929,582158 | -0,726762941 | 0,094945 | -7,65457 | 1,94E-14 | 7,86E-13 |
| Syp | 18196,25343 | -0,300121828 | 0,094431 | -3,17823 | 0,001482 | 0,005261 |
| Flna | 986,3396728 | -0,655434009 | 0,145296 | -4,51101 | 6,45E-06 | 4,52E-05 |
| Gnas | 12692,92313 | -0,578089262 | 0,093203 | -6,20246 | 5,56E-10 | 1,01E-08 |
| Grk2 | 1720,03957 | -0,928316376 | 0,126954 | -7,3122 | 2,63E-13 | 8,66E-12 |
| *Negative values: downregulation in stressed KSCL group* | | | | | | |
| **Comparison Stressed vs Naive** | | | | | | |
| none of the genes above are significantly affected | | | | | | |

__Figure S2:__ **Table S1 showing overlap in significantly upregulated or downregulated mRNAs within the prefrontal cortex of mice exposed to 5d-FSS paradigm associated with AD treatments**. Note that among opioid system genes, only the upregulation of Oprk1, Oprm1, Pdyn and Penk was associated the AD response. **Table 2 shows corresponding values for genes listed in Gene Ontology annotations: "Opiod receptor binding" and "opioid receptor signaling pathway".**
